# A novel fluorobenzothiazole RBx 10080758 as a dual inhibitor against bacterial GyraseB (GyrB) and Topoisomerase IV (parE) of Gram-positive pathogens causing skin and respiratory infections

**DOI:** 10.1101/2022.09.08.507076

**Authors:** Tarani Kanta Barman, Manoj Kumar, Tridib Chaira, Smita Singhal, Tarun Mathur, Vandana Kalia, Ramkumar Gangadharan, Madhvi Rao, Manisha Pandya, Pragya Bhateja, Ruchi Sood, Dilip J. Upadhyay, Shibu Varughese, Ajay Yadav, Lalima Sharma, Naresh Kumar, Jitendra Sattigeri, Pradip K. Bhatnagar, V. Samuel Raj

**Author notes:** Address for Correspondence: Prof. V. Samuel Raj, Centre for Drug Design Discovery and Development (C4D), SRM University, Delhi NCR, Rajiv Gandhi Education City, Sonepat, Haryana – 131029, India.

## Abstract

Development of a novel inhibitor targeting Gyrase B and Topoisomerase IV offers a potential opportunity to combat the drug resistance. In the present study, we extensively investigated the efficacy of RBx 10080758, a novel fluorobenzothiazole, against skin and respiratory infections caused by Staphylococci and Streptococci in *in vitro* and *in vivo* models. RBx 10080758 showed a potent IC_50_ of 0.06 μM against Gyrase and Topoisomerase IV and also exhibited a strong whole cell *in vitro* activity with MIC ranges of 0.015-0.06, 0.015-0.03, 0.008-0.03, 0.008-0.03, 0.008-0.0.06, 0.015-0.06 μg/ml against *Staphylococcus. aureus, Streptococcus pneumoniae*, Coagulase negative Staphylococci, *Streptococcus viridans, Streptococcus pyogenes* and Enterococcus, respectively. As expected from the novel class of molecule, it retains potent even against linezolid and vancomycin resistant strains. Interesting, in mouse systemic infection 10 mg/kg, IV dose protected 100% mice from lethal infections. In rat thigh infection model with MRSA WCUH29 at 45 mg/kg exhibited >3-log_10_ cfu reduction in thigh muscles. RBx 10080758 displayed potent *in vitro* and *in vivo* activity against a panel of MDR Gram-positive bacteria. As a novel chemical class, the fluorobenzothiazoles have the potential to become clinically viable antibiotics, to address the drug resistance problem by its unique dual targeting mechanism of action.

## Introduction

Gram-positive bacteria, particularly nosocomial methicillin resistant *Staphylococcus aureus* (MRSA), have emerged as a superbug, showing resistance to multiple antibiotics [1, 2]. MRSA infection constitutes the most important cause of health care associated infections, and capable of causing various illness ranging from mild skin infection to life-threatening pneumonia and blood stream infection [3, 4]. The treatment for MRSA is quite complicated and expensive as the infections can recur frequently and can easily progress to life-threatening septicemia or atopic dermatitis [5, 6]. The standard of care (SOC) drug of MRSA such as vancomycin, daptomycin, linezolid and tedizolid are promising antibiotics but not completely devoid of limitations [7, 8]. For example, the extended use of linezolid may cause myelosuppression, while tedizolid shows poor activity in neutropenic patients [7]. Daptomycin has poor activity in pulmonary surfactant and also cause myopathy and neuropathy [9]. Therefore, there is a high unmet medical need for a new class of anti-MRSA antibiotic to overcome the limitations of SOC and fight against emerging resistance.

Development of novel inhibitors against DNA Gyrase B (GyrB) and Topoisomerase IV offers a potential opportunity to meet the challenge of resistance [10, 11]. GyrB and Topoisomerase IV are essential enzymes and play important role in DNA replication and compaction. DNA supercoiling activity is essential in all bacteria but not found in human and hence it is an ideal target [12]. The overall identity between *GyrB* and Topoisomerase IV *parE* is 52%, whereas the identity between *GyrB* and *GyrA* is only 5% in MRSA. In addition, identity of bacterial *GyrB* with human topoisomerase is only 12%, which further reduce the possibility of toxicity [13, 14]. The coumarin class of compounds novobiocin and coumermycin bind to GyrB and parE and both target ATP binding sites [15]. In addition, Cyclothialidines are potent inhibitors of Gyr B or GyrB/par E. However, their uses have been limited by poor PK profiles [15, 16].

Therefore, it is of great opportunity to design a dual inhibitor against GyrB and parE, especially directed at the ATP binding sites. Additionally dual-target inhibitors will pose lower risks of resistance development than single-target inhibitors. RBx 10080758 is a novel fluorobenzothiazole inhibits both GyrB and Topoisomerase IV parE of Gram-positive bacteria. In present study, the *in vitro* and *in vivo* activities of RBx 10080758 with pharmacokinetics against Gram positive bacteria particularly MRSA were evaluated.

## Results

### Enzyme inhibitory activity of RBx 10080758 and RBx 10115912

RBx 10080758 and RBx 10115912 (figure 1) showed a strong inhibitor activity against bacterial gyrase and topoisomerase IV. The 50% inhibitory concentrations (IC_50s_) of RBx 10080758 and RBx 10115912 for *E. coli* gyrase supercoiling and topoisomerase IV decatenation assay are shown in Table 1. Both compounds showed no inhibitory activity against human topoisomerase II up to 500 μM (Table 1), suggesting their high selectivity to bacterial topoisomerases.

**Figure 1.**
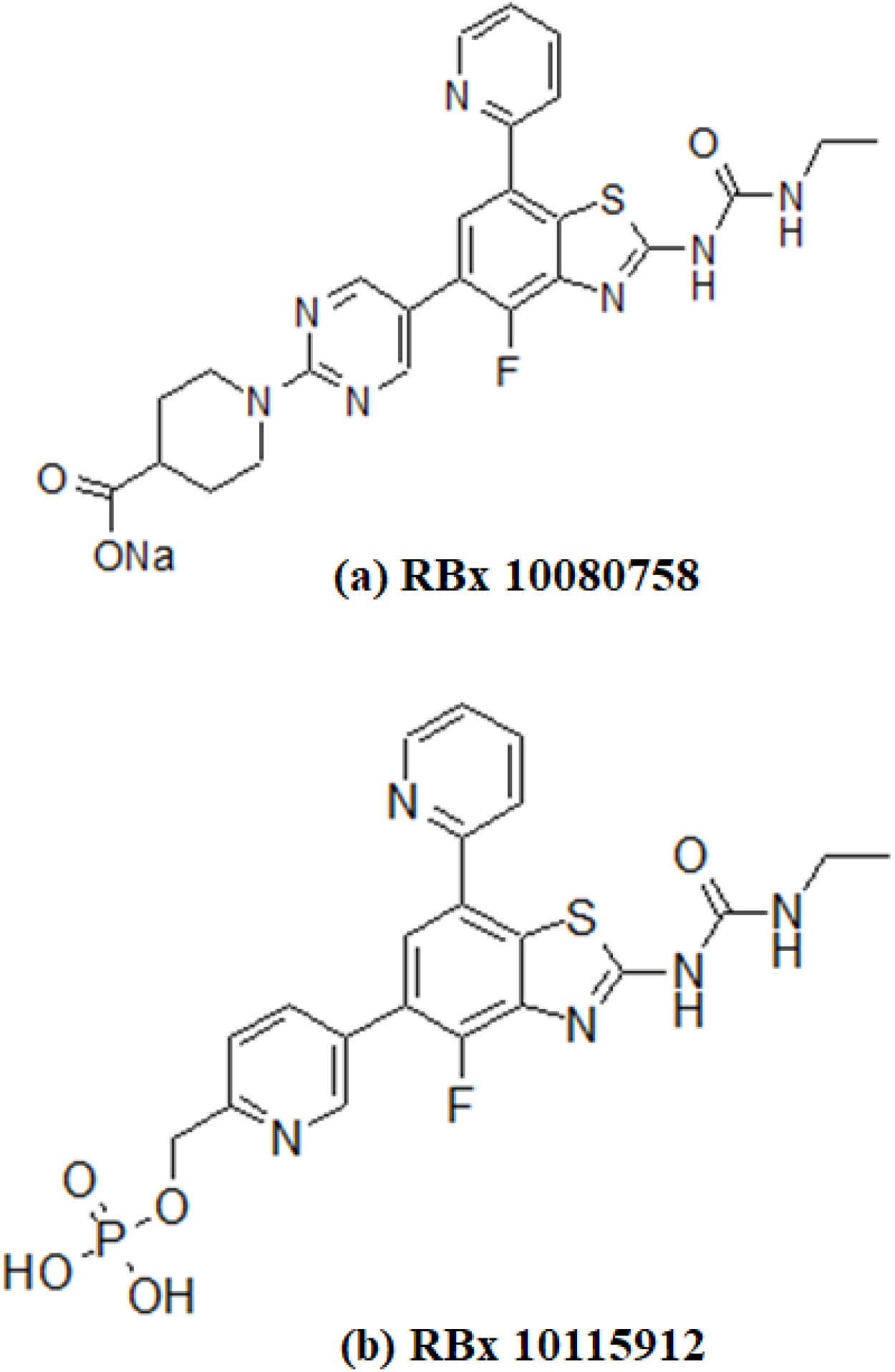
Chemical structure of RBx 10080758(a) and RBx 10115912(b)

**Table 1:** IC_50_ of RBx 10080758 and RBx 10115912 against *E. coli* DNA gyrase, Topoisomerase IV and human Topoisomerase II

| Compound | IC<sub>50</sub> (μg/ml)<sup>a</sup> |  |  |
| --- | --- | --- | --- |
|  | *E. coli* |  | Human |
|  | DNA Gyrase | Topoisomerase IV | Topoisomerase II |
| RBx 10080758 | 0.06 | 0.06 | >500 |
| RBx 10115912 | 0.06 | 0.06 | >500 |
| Levofloxacin | 54 | 7.5 | >500 |

<sup>a</sup>All reactions were performed in duplicate and repeated twice. The mean values were used to determine the IC<sub>50</sub>.

### Antibacterial activity against Gram-positive bacteria

The *in vitro* activities of RBx 10080758 and RBx 10115912 against a panel of Gram-positive bacteria such as *S. aureus, S. pneumoniae*, coagulase negative Staphylococci, *S. viridians, S. pyogenes and Enterococcus* are summarized in Table 2. RBx 10080758 displayed stronger inhibitory activity against recent clinical isolates of *S. aureus* including MSSA, MRSA (n=50) and multiple drug resistant (MDR) pathogens with MIC_90_ of 0.06 μg/ml and MIC range of 0.015-0.06 μg/ml (Table 2). Overall, both RBx 10080758 and RBx 10115912 displayed potent activity against multiple MDR bacteria (Supplementary figure 1a-d).

**Table 2:** *In vitro* activity of RBx 10080758 and RBx 10115912 in comparison to other antimicrobial agents against Gram positive clinical isolates

| Organisms (no of strains)/new chemical entities/standard drugs | MIC (µg/ml) |  |  |
| --- | --- | --- | --- |
|  | MIC<sub>50</sub> | MIC<sub>90</sub> | MIC Range |
| ***Staphylococcus aureus* (n=50)<sup>1</sup>** |  |  |  |
| RBx 10080758 | 0.03 | 0.06 | 0.015-0.06 |
| RBx 10115912 | 0.25 | 0.5 | 0.125 - 1 |
| Linezolid | 2 | 2 | 1-2 |
| Vancomycin | 0.5 | 1 | 0.5 - 1 |
| Ciprofloxacin | 0.5 | >16 | 0.25->16 |
| Erythromycin | 0.5 | >16 | 0.25->16 |
| Mupirocin | 0.25 | 256 | <0.125-256 |
| ***Streptococcus pneumoniae* (n=20)** |  |  |  |
| RBx 10080758 | 0.015 | 0.03 | 0.015-0.03 |
| RBx 10115912 | 0.125 | 0.25 | 0.125–0.5 |
| Linezolid | 1 | 2 | 0.5- 1 |
| Vancomycin | 0.5 | 1 | 0.5–1 |
| Ciprofloxacin | 0.5 | 2 | 0.5-4 |
| Erythromycin | 0.06 | 16 | 0.06->16 |
| Penicillin | 0.03 | 0.25 | 0.03->4 |
| **Coagulase negative Staphylococci (n=35)<sup>2</sup>** |  |  |  |
| RBx 10080758 | 0.015 | 0.03 | 0.008-0.03 |
| RBx 10115912 | 0.25 | 0.5 | 0.125 - 1 |
| Linezolid | 1 | 1 | 0.5-1 |
| Vancomycin | 2 | 2 | 1-2 |
| Ciprofloxacin | 0.5 | >16 | 0.125->16 |
| Erythromycin | 0.25 | >16 | 0.125->16 |
| ***Streptococcus viridans* group (n=20)** |  |  |  |
| RBx 10080758 | 0.008 | 0.03 | 0.008-0.03 |
| RBx 10115912 | 0.25 | 0.5 | 0.125 - 4 |
| Clindamycin | 0.03 | 0.06 | 0.03-0.06 |
| Amoxicillin | 0.06 | 0.25 | 0.03-0.25 |
| Erythromycin | 0.03 | 0.06 | <0.015-0.06 |
| ***Streptococcus pyogenes* (n=50)<sup>3</sup>** |  |  |  |
| RBx 10080758 | 0.03 | 0.03 | 0.008-0.06 |
| RBx 10115912 | 0.25 | 0.5 | 0.125 - 2 |
| Linezolid | 1 | 2 | 0.5-2 |
| Erythromycin | 0.03 | >4 | 0.015->4 |
| Penicillin | 0.008 | 0.008 | <0.004-0.008 |
| Ciprofloxacin | 0.5 | 4 | 0.5-4 |
| Clindamycin | 0.03 | 4 | 0.015->4 |
| **Enterococci spp (n=27)<sup>4</sup>** |  |  |  |
| RBx 10080758 | 0.015 | 0.06 | 0.015-0.06 |
| RBx 10115912 | 0.25 | 0.5 | 0.125 - 2 |
| Linezolid | 2 | 4 | 1-4 |
| Vancomycin | 1 | >16 | 1>16 |
| Amoxicillin | 0.5 | >16 | 0.25->16 |
| ***S. aureus* LZD<sup>R</sup> (n= 10)** |  |  |  |
| RBx 10080758 | 0.03 | 0.06 | 0.015-0.06 |
| RBx 10115912 | 0.25 | 0.5 | 0.125 - 1 |
| Linezolid | 32 | >32 | 8->32 |
| Vancomycin | 1 | 2 | 0.5-2 |
| Ciprofloxacin | 1 | >16 | 0.25->16 |
| Erythromycin | 2 | 4 | 0.25-8 |
| ***S. aureus* VISA or VRSA strains (n=21)<sup>5</sup>** |  |  |  |
| RBx 10080758 | 0.03 | 0.03 | <0.008-0.03 |
| RBx 10115912 | 0.25 | 0.5 | 0.125 - 1 |
| Linezolid | 2 | 2 | 1-2 |
| Vancomycin | 4 | 4 | 4-16 |
| Ciprofloxacin | 0.5 | 16 | 0.25->16 |
| Erythromycin | >16 | >16 | 0.5 - >16 |

<sup>1</sup>Methicillin sensitive *S. aureus* (MSSA), methicillin resistant *S. aureus* (MRSA), fluoroquinolone resistant (FQ<sup>R</sup>) *S. aureus*, Erythromycin<sup>R</sup> *S. aureus*, Mupirocin<sup>R</sup> *S. aureus*, <sup>2</sup>Methicillin sensitive *S. epidermidis* (MSSE), methicillin resistant *S. epidermidis* (MRSE), *S. haemolyticus* & other coagulase negative staphylococcus (Cons), 3. Sensitive, erm & mef strains, 4. *E. faecalis* and *E. faecium*, 5. Vancomycin intermediate *S. aureus* (VISA, n=19) and Vancomycin resistant *S. aureus* (VRSA, n=2)

### *In vitro* cytotoxicity

Cytotoxicity assessment of RBx 10080758 and RBx 10115912 was performed using standard MTT assay. The IC_50_ values were calculated using Graph Pad Prism and both compounds showed IC_50_ value of ≥50 μg/ml.

### Frequency of resistance

Though RBx 10080758 and RBx 10115912 showed potent activity against the panel of bacterial pathogens, RBx 10115912 showed poor solubility hence was not evaluated for further studies. The frequency of spontaneous resistance (FSR) development was determined at 4 × and 8 × the MIC of RBx 10080758 against MRSA. The FSR at 4 × MICs of RBx 10080758 was <3.5 × 10^-9^ against MRSA WCUH-29 and <6 × 10^-9^ against MRSA 562 (Table 3). RBx 10080758 showed mutation prevention concentration (MPC) of 0.125 μg/ml and completely prevented the spontaneous resistant development at 0.125 μg/ml.

**Table 3:** Frequency of resistance and MPC RBx 10080758 against *S. aureus*

| Bacterial Strains | Frequency of Resistance |  | MIC (µg/ml) | MPC | MPC/MIC ratio |
| --- | --- | --- | --- | --- | --- |
|  | 4 × MIC | 8 × MIC |  |  |  |
| *S. aureus* WCU H 29 | <3.5 ×10<sup>-9</sup> | <3.5 ×10<sup>-9</sup> | 0.03 | 0.125 | 4 |
| *S. aureus* MRSA 562 | <6.0 ×10<sup>-9</sup> | <6.0 ×10<sup>-9</sup> | 0.03 | 0.125 | 4 |

### Combination studies of RBx 10080758

RBx 10080758 was checked for activity in combination with ten existing drugs against *S. aureus* strains. The FIC index against the antibiotics are as follows: Linezolid (FIC index 1), Vancomycin (1.5), Gentamicin (1), Ciprofloxacin (1), Erythromycin (2), Ertapenem (1), Imipenem (1.2), Augmentin (1.5), Clindamycin (1.5) and Cefoxitin (1.25). The FIC index of RBx 10080758 against all the antibiotics is indicative of an additive effect (data not shown). RBx 10080758 shows no synergy or antagonism with existing drugs being used for treatment against *S. aureus*.

### Time-kill kinetic studies

The results of time-kill studies of RBx 10080758 and linezolid against MRSA 562, MRSA WCUH-29, *S. pneumoniae* MA 80 and *S. pyogenes* are presented in Figure 2 a-d. RBx 10080758 showed rapid concentration dependent killing of MRSA isolates (Figure 2a, 2b). However, RBx 10080758 exhibited time-dependent bactericidal activity against *S. pneumoniae* MA-80 and *S. pyogenes* even at 1 × MIC (Figure 2c, 2d).

**Figure 2:**
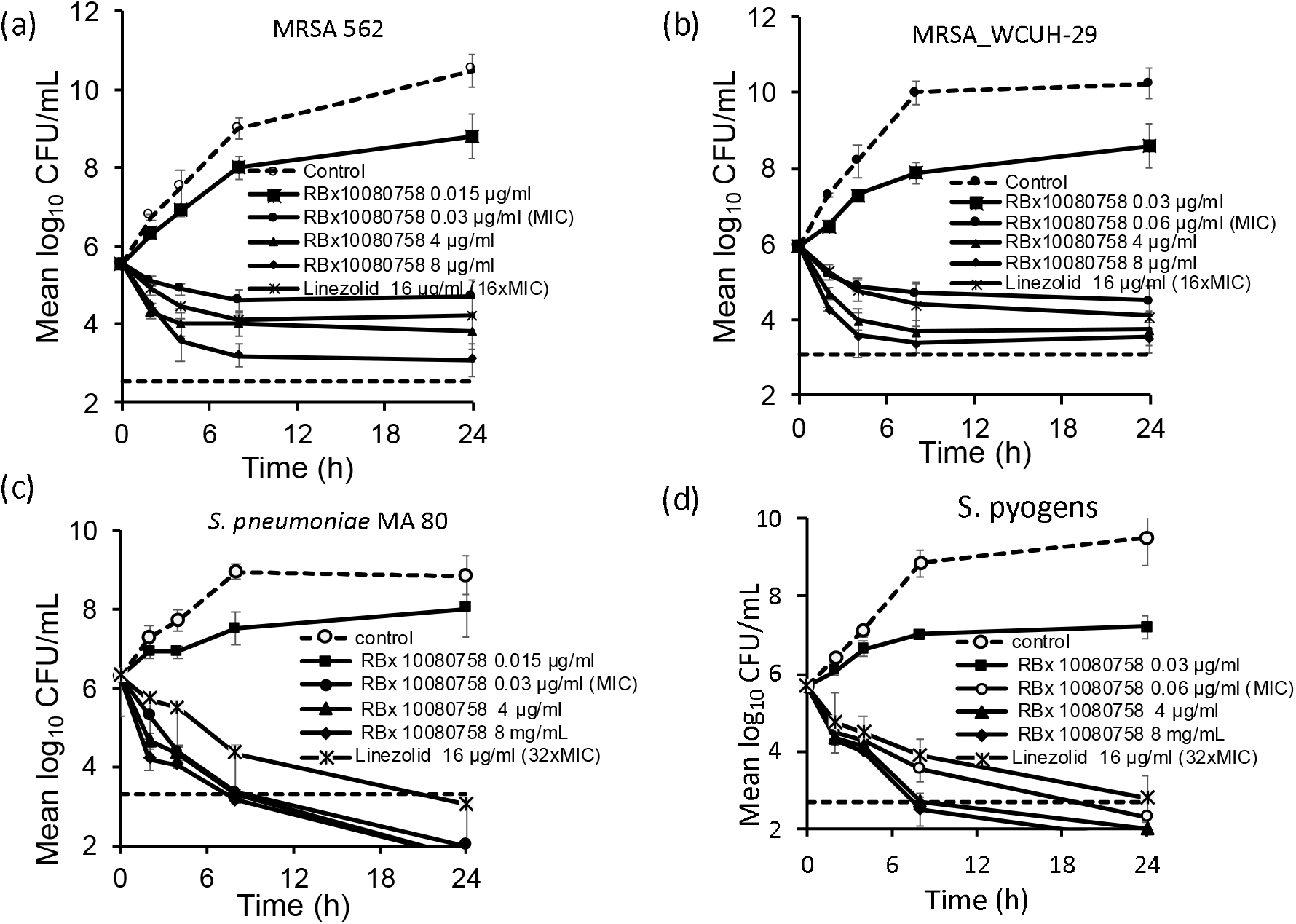
Time-kill curves of RBx 10080758 against MRSA 562 (a), MRSA WCUH-29 (b), *S. pneumoniae* MA-80 (c) and *S. pyogenes* (d). Data represent mean ± SD from three independent experiments.

### Murine systemic infection model of *MRSA-562*

The *in vivo* efficacy of RBx 100 80758 in murine systemic infection model of MRSA-562 is presented in Figure 3. All the infected mice treated with placebo, started developing the clinical symptoms of infection with 8 h of infection and died by following day. RBx 10080758 showed a dose dependent efficacy against MRSA-562 (Figure 3a) with ED_50_ of 5 mg/kg. The kidney bacterial load at 8 h or 24 h post-infection, showed significant reduction in bacterial count at 10 and 20 mg/kg dose as compared to placebo control or linezolid (Figure 3b), suggesting the superior killing potential of RBx 10080758.

**Figure 3:**
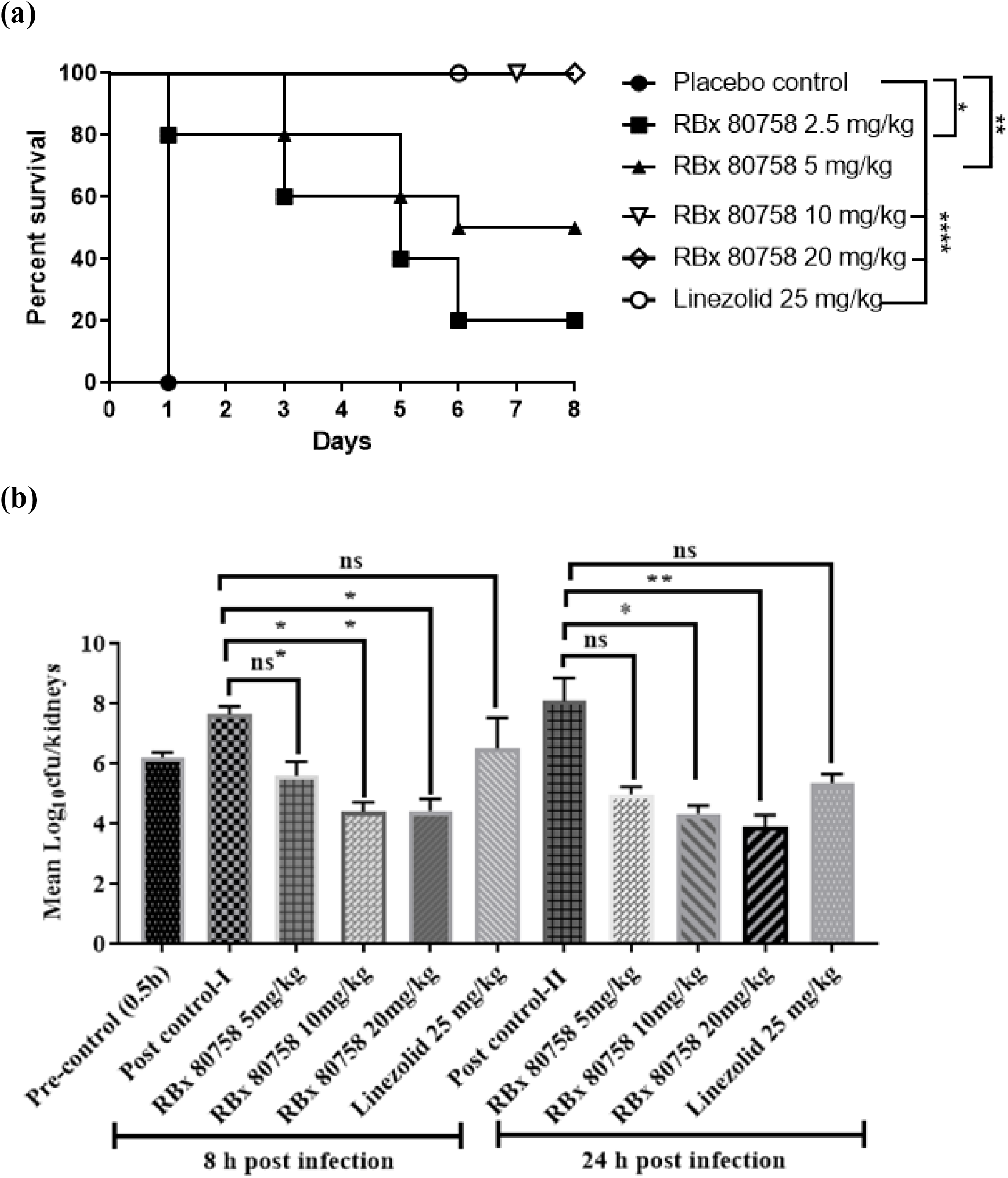
Efficacy of RBx 10080758 in murine systemic infection model of MRSA-562. (a) Kaplan Meier survival curve (n=6-10), (b) Bacterial load in the kidneys mouse (n=4) treated with RBx 10080758 at 8 h after two doses and at 24 h after 4 four doses. Asterisks indicate a significant difference compared with the result of the post-control group. Survival data were analyzed by log-rank Mantel-Cox test. Kidney bacterial loads were compared using one-way analysis of variance (ANOVA) with Dunnett’s multiple-comparison posttest. (\**P* < 0.05, \*\**P* < 0.01, \*\*\**P* < 0.001, \*\*\*\**P* <0.0001).

### Rat thigh and lung infection models

The efficacy of RBx 10080758 (Na salts) was investigated against MRSA WCUH-29 in rat thigh infection model (Fig. 4a–4c). RBx 10080758 showed a dose dependent efficacy against MRSA WCUH-29 with a bactericidal potential at 45 mg/kg, IV dose (Figure 4a). When tested against the highly virulent MRSA PVL-2 strain, RBx 10080758 exhibited similar bactericidal efficacy at 45 mg/kg dose (Figure 4b). Similar activity of RBx 10080758 was also observed against MRSA WCUH-29 at 45 mg/kg dose in 2 days infection model (Figure 4c).

**Figure 4:**
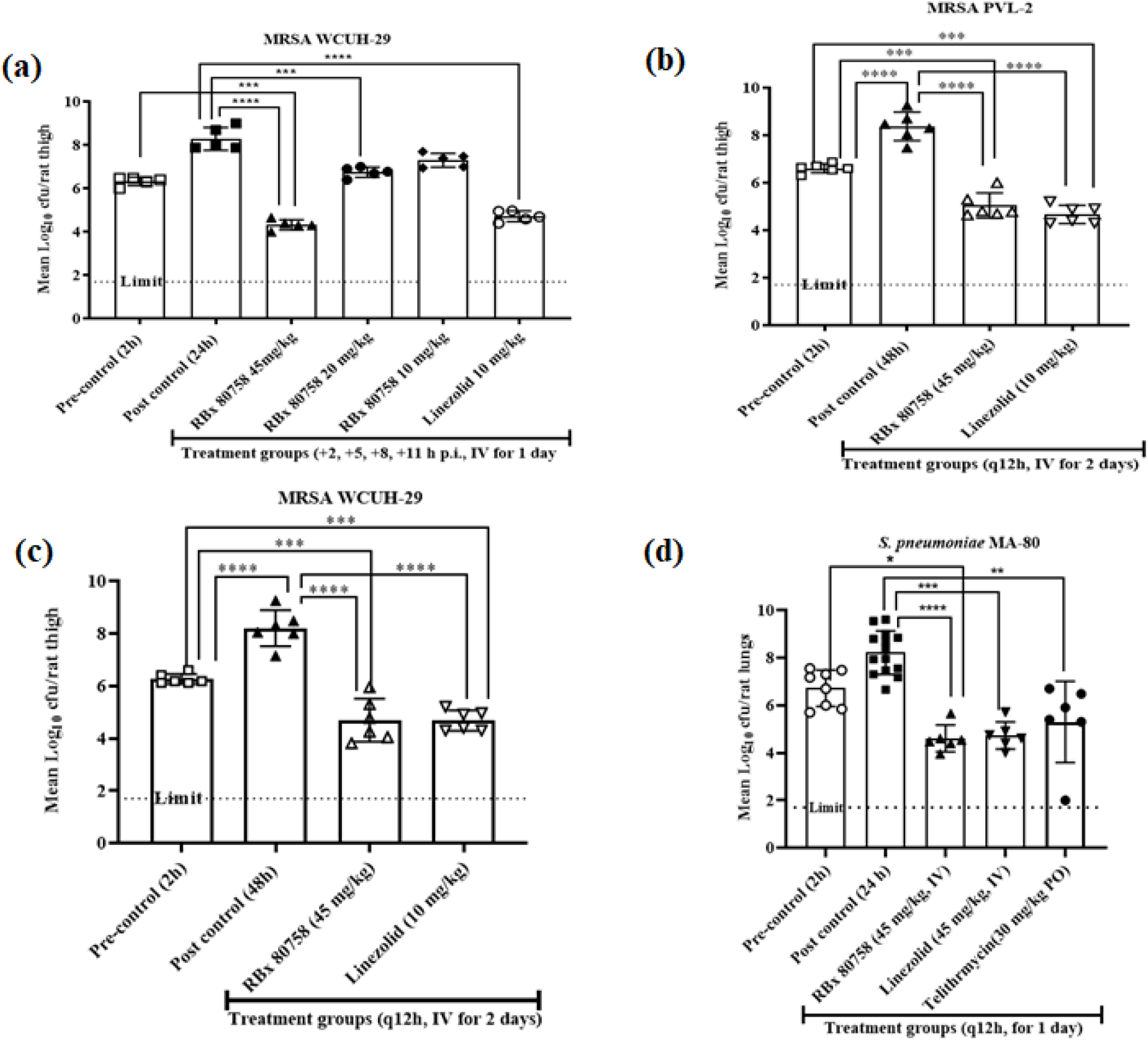
Efficacy of RBx 10080758 (Na salts) in rat thigh and lung infection models. Each bar represents the mean ± standard deviation (n=5-14). Rat thigh infection was performed with MRSA WCUH-29 with four times a day dosing regimen for 1 day treatment (**a**); MRSA PVL-2 with q12h x 2 days (**b**) and MRSA WCUH-29 with q12h x 2 days treatment (**c**); or intratracheal infection with *S. pneumoniae* MA-80 with q12h x 1 day (**d**). Treatment was initiated 2 h post-infection. Bacterial counts in each treated group were compared with those of pre- and post-control animals. The asterisks indicate a significant difference in bacterial counts after treatment compared to pre-and post-control (\**P*< 0.05, \*\**P*< 0.01, \*\*\**P*< 0.001, \*\*\*\**P* <0.0001).

In the rat lung infection of *S. pneumoniae* MA 80, RBx 10080758 exhibited a bactericidal effect at 45 mg/kg dose with a significant >3 log_10_ reduction in bacterial load as compared to placebo control. Linezolid and telithromycin also exhibited a similar effect against *S. pneumoniae* MA-80 (Figure 4d).

### *In vitro* clearance RBx 10080758 in liver microsomes

The clearance of RBx 10080758 in human liver microsomes contributed by glucuronidation was assessed. The results revealed a minimal involvement of Phase II glucuronidation in the clearance of RBx 10080758. The clearance of RBx 10080758 in liver microsomes was stable and this is not a high clearance molecule.

In CYP inhibition assay, all the compounds from 4-fluorobenzothiazole series (data not shown) have shown IC_50_ for 5 CYP enzymes (CYP1A2, CYP2C9, CYP2C19, CYP2D6 and CYP3A4) >10 μM concentration. Intrinsic clearance of 4-fluorobenzthiazole/pyrrole ABH amide series of compounds are metabolically stable, and they have shown low-moderate clearance in rat and human liver microsomes (<0.6 mL/min/kg). RBx 10080758 and RBx 10115912 are two representative compounds of these series.

### *In Vivo* pharmacokinetics of RBx 10080758

*In vivo* pharmacokinetic data of RBx 10080758 are presented in Table 4. RBx 10080758 displayed similar intravenous pharmacokinetics in rat and dog. In both the species, it showed low mean plasma clearance (rat 11.34 ml/min/kg and dog 13.67 ml/min/kg), low V_ss_ (< 0. 7) and short half-life (< 2.0 h). The plasma exposure (AUC_inf_) of RBx 10080758 was 0.07, 1.5 and 1.27 μg.h/ml in mouse, rat, and dog, respectively by IV route.

**Table 4:** Pharmacokinetics of RBx 10080758 (0-8 h) in rat and dog by intravenous route

| Species | IV Dose (mg/kg) | C<sub>max</sub> (µg/ml) | t<sub>1/2</sub> (h) | AUC<sub>inf</sub> (µg.h/ml) | CL<sub>p</sub> (ml/min/kg) | V<sub>ss</sub> (l/kg) |
| --- | --- | --- | --- | --- | --- | --- |
| Rat | 1 | 5.19 ± 0.50 | 1.63 ± 0.50 | 1.50 ± 0.52 | 11.34 ± 0.49 | 0.17 ± 0.03 |
| Dog | 1 | 4.38 ± 1.63 | 1.14 ± 0.51 | 1.27 ± 0.30 | 13.67 ± 3.39 | 0.39 ± 0.12 |

## Discussion

RBx 10080758 exhibited potent activity against both *S. aureus* DNA GyrB and Topoisomerase IV enzymes and showed >1000-fold selectivity against human Topoisomerase II. Usually, gyrase inhibitors are more active against DNA gyrase than topoisomerase, while quinolones show more inhibitory potency against Topoisomerase IV than that of GyrB [17, 18]. Interestingly, RBx 10080758 inhibited *S. aureus* DNA GyrB and Topoisomerase IV with equal potency, which may significantly reduce the risk of resistance development. As RBx 10080758 is a novel dual inhibitor of DNA GyrB and Topoisomerase IV and exhibited potent MIC against various Gram-positive bacteria with no cross-resistance to fluoroquinolones or other antibiotics, which highlights its utility against MDR pathogens. RBx 10080758 showed excellent *in vitro* activity against a panel of skin and soft tissue and respiratory pathogens. RBx 10080758 showed ≥16-fold potent MIC as compared to ciprofloxacin and ≥60-fold potent MIC as compared to linezolid against *S. aureus*. Most clinical isolates of MRSA are resistant to fluoroquinolones, which limits their clinical utilities [17, 19]. In addition, emergence of large number of MDR gram-positive strains in nosocomial settings even against recently developed linezolid and tedizolid, limiting the treatment options against these MDR pathogens [20]. Although, in the recent years, several GyrB and ParE inhibitors demonstrated potent *in vitro* activities against MDR Gram-Positive pathogens, however none have showed acceptable *in vivo* efficacy or progress for clinical development.

Interestingly, RBx 10080758 exhibited superior activities against the different MDR strains such as MDR *S. aureus*, linezolid resistant (LZD^res^) *S. aureus*, vancomycin-intermediate *S. aureus* (VISA), vancomycin resistant *S. aureus* (VRSA), vancomycin resistant Enterococci (VRE), LZD^res^ *S. pyogenes*, fluoroquinolone resistant (FQ^res^) *S. viridians* and *S. pneumoniae* (Supplementary figure 1a-d) and showed low frequency of spontaneous resistance, suggest that RBx 10080758 may effectively cover both susceptible and MDR pathogens and will have a low propensity for resistance development in clinical settings [21]. RBx 10080758 exhibited low MPC of 0.125 μg/ml which may be useful in selecting appropriate dosing of drug for efficacy studies and determining the potential of drug in preventing the selection of resistant mutants. Pulmonary surfactant reduces the surface tension at the air/liquid interface of alveoli and is known to significantly impair the antimicrobial activities of moxifloxacin, colistin and daptomycin [9, 12, 22, 23]. However, the activity of RBx 10080758 was not affected by the presence of a lung surfactant (Supplementary Figure 2), demonstrating its effectiveness in pulmonary infections.

Though, RBx 10080758 was found metabolically stable, however, its poor solubility hindered the further evaluation. Different salt forms of RBx 10080758 such as sodium, lithium and hydrochloric acid salts were explored to improve its solubility; sodium salt of RBx 10080758 showed significantly better solubility, stability, and exhibited the similar MIC activity against bacterial pathogens (data not shown). Hence sodium salt of RBx 10080758 was selected for *in vivo* evaluation. To confirm the broad-spectrum gram-positive pathogen coverage, the efficacy of RBx 10080758 was assessed, in rat thigh model against *S. aureus* and in lung infection against *S. pneumoniae* as well as mouse systemic infection models caused by MRSA. These models bear a resemblance to human skin and soft tissue infection (SSTI), pneumonia and septicemia. RBx 10080758 showed excellent *in vivo* efficacy with ≥3 log_10_ reduction in one day as well as 2 days rat thigh infection model against *S. aureus* indicating its suitability for SSTI indications. RBx 10080758 also showed potent efficacy against *S. pneumoniae* in rat lung infection model, indicating the potential for lung infection.

In systemic infection rapid clearance of pathogen is extremely important to control the spread of infection. RBx 10080758 showed 100 % survival at 10 mg/kg dose in murine systemic infection model of MRSA-562 and exhibited a significant reduction in the kidney bacterial loads as compared to linezolid and untreated control. This indicated superior clinical and microbiological cure by RBx 10080758 compared to linezolid. To calculate the clinical efficacious dose of RBx 10080758 and to confirm safety, further animal studies are warranted [24, 25]. RBx 10080758 with low clearance and low volume of distribution, yielded AUC_inf_ (μg.h/ml) and C_max_ that correlated well with *in vivo* efficacy in rat. There is likely an improvement in efficacy in dog (non-rodent species) due to similar pharmacokinetics observed between rat and dog. Taken together, RBx 10080758 exhibited potent *in vitro* and *in vivo* activities against MRSA and *S. pneumoniae*. These results indicate that RBx 10080758 can be explored as a new therapeutic agent against Gram-positive bacterial infections that cause SSTI and pneumonia.

## Materials and Methods

### Bacterial strains, antibiotics and new chemical entities

Different bacterial isolates, such as, *Staphylococcus aureus* (n=81) including MSSA, MRSA, vancomycin and linezolid resistant strains, *Staphylococcus epidermidis* (n=35), *Streptococcus viridans* group (n=20), Enterococci (n=27) including *Enterococcus faecalis* and *Enterococcus faecium* and *Streptococcus pyogenes* (n=50) were used in this study. Standard strains were purchased from the American Type Culture Collection (ATCC, Manassas, VA, USA), and clinical isolates were acquired from Indian hospitals. RBx 10080758 (Figure 1a) and RBx 10115912 (Figure 1b) were synthesized in-house at Daiichi Sankyo India Pharma Pvt. Ltd., Gurgaon, India. Standard antibiotics were purchased from commercial sources.

### DNA Gyrase supercoiling assay

*E.coli* DNA gyrase supercoiling assay was performed as we described earlier [18]. Briefly, the assay was carried out in a reaction volume of 20 μl containing 1 unit of DNA gyrase and 0.5 μg of relaxed pBR322 DNA in assay buffer in presence of RBx 10080758, RBx 10115912 or levofloxacin. After incubation at 37°C for 60 min with and without inhibitors, the reaction was stopped. The amount of the supercoiled DNA band was quantified using Bio-Rad’s Quantity one software.

### Topoisomerase IV decatenation assay

DNA decatenation assay was performed using *E. coli* Topoisomerase IV (subunits A and B) as described earlier [11, 18]. The assay was performed in 20 μl of a reaction mixture containing 1 unit of Topoisomerase IV and 0.4 μg of catenated kinetoplast pBR322 DNA in assay buffer in presence or absence of test compounds. After incubation at 37°C for 60 min, the reaction was stopped. The IC_50_ values were determined as described earlier [18].

### Activity against human topoisomerase IIα

Human Topoisomerase-IIα was synthesized as a histidine-tagged protein in *E. coli*.Decatenation assays were performed in 20 μl reaction mixture which include 50 mM Tris-HCl (pH 7.5), 120 mM KCl, 10 mM MgCl_2_, 0.5 mM ATP, 0.5 mM DTT, 30 μg/ml BSA, 0.4 μg catenated kinetoplast DNA as a DNA substrate, human topoisomerase IIα enzyme, and test compounds. After incubation at 37 °C for 60 min, the reaction was terminated and IC_50_s were determined as described above. All the above experiment was performed in duplicate and repeated twice.

### *In vitro* susceptibility testing

Minimum inhibitory concentrations (MIC) of RBx 10080758, RBx 1011591, and other standard antibiotics were determined against a panel of Gram-positive bacterial strains using micro-broth dilution method as recommended by the Clinical and Laboratory Standards Institute (CLSI) [26]. All experiments were performed in triplicate, and highest MIC values were considered as the final results. To determine the effect of pulmonary surfactant on *in vitro* activity of RBx 10080758, RBx 1011591, the MIC was also determined in presence of pulmonary surfactant as described earlier [9]

### *In vitro* cytotoxicity assay

Cytotoxicity assay was conducted using MTT assay. HepG2 cells were seeded in a 96-well flat-bottom microtiter plate and allowed to adhere for 24 h at 37°C in a CO_2_ incubator. After 24 h of incubation, cells were treated with 200 μl of the test compound. After 48 h of incubation, 12.5 μl of MTT working solution (5 mg/ml) was added and incubated again for 4 h. Finally, the intensity of the DMSO dissolved formazan crystals was quantified at 540 nm.

### Combination study of RBx 10080758

Checkerboard assay was performed to evaluate their fractional inhibitory concentration (FIC) index in combinations against standard MRSA strain, and the assay were performed on 96-well plates as described previously [27, 28]. For example, combination A/B was used for interpreting the assay protocol [29]. The FICs were calculated as described earlier. The combining effect of A in combination with B against MRSA was interpreted as described earlier [28–30].

### Time-kill study

The time-kill studies of RBx 10080758, linezolid, against MRSA-562, MRSA WCUH-29, *S. pneumoniae* MA-80 and *S. pyogenes* were performed as described earlier [22, 31]. In brief, cation-adjusted Mueller-Hinton broth (CA-MHB) was pre-warmed at 37°C, and the drug was added at concentrations ranging from 0.5 × MIC to 8 μg/ml. Then the exponential-phase cultures were added to the culture flask (~5 × 10^6^ cfu/ml). For *S. pyogenes* and *S. pneumoniae*, CA-MHB with 5% lysed horse blood and drug were inoculated with ~5 × 10^6^ cfu/ml, and the flasks were incubated at 35°C. The viable cell count was assessed at predefined time points. The time-kill kinetics were performed three independent times.

### Frequency of resistance

The frequency of resistance was determined as described earlier [18, 32]. Bacterial inoculums of each *S. aureus* strain were prepared in MHB, and 100 μl of each suspension was spread onto the surface of MHA containing 4 × and 8 × MIC of RBx 10080758. After incubating for about 48 h at 37°C, the number of colonies was counted and divided by the number of bacteria inoculated to determine the frequency of resistance.

### *In vitro* ADME studies

For determination of intrinsic clearance in liver microsomes, the reaction mixture was prepared by addition of 400 μl of NADPH regenerating solution, 25 μl of liver microsomes and made up to volume of 995 μl with phosphate buffer (100 mM, pH 7.4). Then 5 μl (100 μM stock) of test compounds was added to initiate the metabolic reaction at 37°C. Periodically 70 μl of aliquots were withdrawn every 3 min for liquid chromatography-tandem mass spectrometry (LC-MS/MS) estimation. The remaining parent compound was estimated at each time point from the samples and was expressed as percentage of 0 min. The intrinsic clearance was expressed in units of L/h/kg.

*In vitro* CYP inhibition assay was performed using a standard commercial recombinant CYPs kit (BD-Gentest) as recommended by the company. Seven serial half-log dilutions of RBx 10080758 and standard inhibitor starting from 100 μM were prepared. The reaction was initiated by mixing of 100 μl of test compound/standard inhibitor with 100 μl of enzyme substrate mix. After incubation, fluorescence was read using a plate reader and IC_50_ was calculated.

### Experimental animals and ethical approval

Specific pathogen free (SPF) and Sprague Dawley (SD) rats were used for efficacy studies, while SPF SD rats and Beagle dogs were used for PK studies. The laboratory environmental conditions were temperature: 22 ± 2°C; relative humidity: 40 to 60 % and day-light cycle: 12-h light/12-h dark. All animal protocols (IAEC/2010/38, IAEC/2010/42, DS/IAEC-2012/006 and IAEC/2010/77/08-E2/10) were approved by the Institutional Animal Ethics Committee.

### Murine systemic infection model of *MRSA-562*

The minimum lethal dose of MRSA for systemic infection was determined as described earlier [33, 34]. The *in vivo* efficacy of RBx 10080758 was determined in murine systemic infection caused by MRSA 562 strain. The bacterial strain was grown in trypticase soy broth at 37°C. The culture was mixed with equal volume of 5% hog mucin (Becton, Dickinson and Company) and 0.5 ml of the bacterial suspension (1-5 × 10^6^ cfu/mouse, n=6-10/group) was injected by intraperitoneal (IP) route into each mouse. RBx 10080758 was administered 2.5, 5, 10, and 20 mg/kg by IP route at +0.5, +4, +7, +10 h post infection (p. i) and linezolid 25 mg/kg was administered orally at +0.5, +6 h p. i. for 1 day. The survival of animals was recorded daily for 8 days.

### Organ load model

Additionally, to assess the microbiological cure of RBx 10080758, two satellite groups of mice (n=4/group) in similar experimental conditions as described above, were kept and their kidney bacterial loads were determined as described earlier [35]. Bacterial load was determined at 8 h and 24 h p. i. in treatment group that received 5, 10, and 20 mg/kg of RBx10080758 and 25 mg/kg of linezolid.

### Efficacy of RBx 10080758 in SD rat thigh infection

SD rats of either sex weighing 90 ±10 g (n=5-6/group) were used for thigh infection model. The overnight grown bacterial culture was diluted in MHB and 250 μl containing 1-3 × 10 cfu was injected in the thigh muscles of each rat. Treatment was initiated 2 h p. i and a pre-control group was sacrificed, and their thigh muscles were evaluated for initial bacterial loads. The efficacy of RBx 10080758 (sodium salts) was evaluated at 10, 20 and 45 mg/kg (molar equivalent dose of RBx 10080758) four times a day at +2, +5, +8, and +11 h p. i and linezolid was evaluated at 10 mg/kg by IV route. At the end of 24 h, the animals were sacrificed, and their thigh muscle was processed for bacterial counts. In addition, the efficacy of RBx 10080758 at 45 mg/kg, q12h for 2 days by IV route was also evaluated against MRSA PVL-2 and WCUH-29 strains in rat thigh infection model.

### Rat lung infection model

To evaluate the efficacy of RBx 10080758 against respiratory tract pathogens, a rat lung infection model was established using *S. pneumoniae* MA-80. SD rats of either sex weighing 80-100 g (n=6-14) were used for lung infection. The overnight grown culture was suspended in PBS and OD_600nm_ was adjusted to 0.3. The bacterial inoculum was further mixed with an equal volume of 5% mucin. The pre-anesthetized SD rats with isoflurane, were inoculated intra-tracheally with 1-5 × 10^6^ cfu/rat as previously described [36]. The treatment of RBx 10080758 and linezolid was started 2 h p. i. with 45 mg/kg, q12h, IV for 1 day and telithromycin was administered at 30 mg/kg by oral gavage. Following 12 h of last dose, lungs of each animal were harvested aseptically and homogenized in 5 ml of PBS and enumerated the viable bacterial counts.

### *In vivo* pharmacokinetic study

Sprague Dawley rats and Beagle dogs were administered single dose of RBx 10080758 at 1 mg/kg by IV route. Serial blood sampling was carried out in rat and dog at different time points up to 8 h post-dose. Plasma was harvested by centrifugation of blood and kept at −80°C until analysis. RBx 10080758 concentrations in plasma were determined using LC-MS/MS. The pharmacokinetic data were analyzed using the NCA module of WinNonlin software (version 4.1).

### Statistical analysis

All data were analyzed using GraphPad Prism Software (version 5.03; San Diego, CA). The statistical significance between the treated and untreated control group was evaluated by nonparametric Mann-Whitney analysis. The bar chart and scatter line graph presented as Means ± SD, were calculated using Microsoft Office Excel. Survival curves were analyzed by log-rank Mantel-cox test. A *P* value of <0.05 was considered statistically significant.

## Transparency declarations

None to declare

## Acknowledgements

Authors are thankful to Ranbaxy and Daiichi Sankyo India Pharma Private Limited for providing necessary facility to carry out the studies.

