## Supplementary file for "A novel fluorobenzothiazole RBx 10080758 as a dual inhibitor against bacterial GyraseB (GyrB) and Topoisomerase IV (parE) of Gram-positive pathogens causing skin and respiratory infections"

### Supplementary Figure

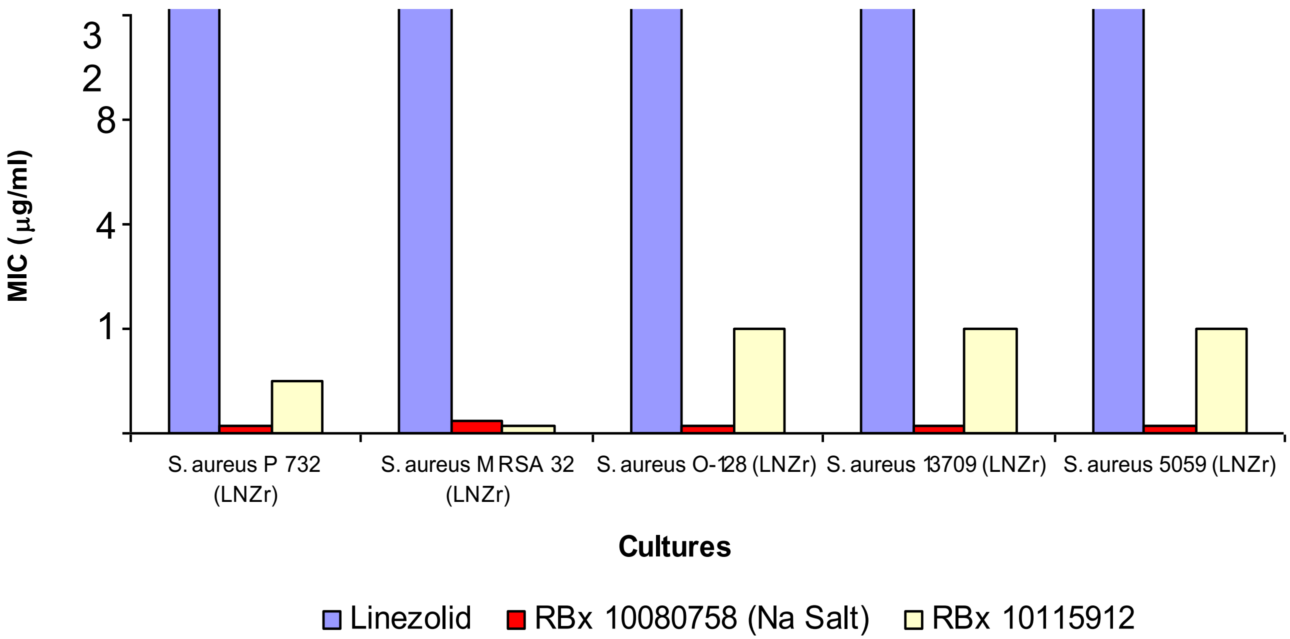


**MIC90 (μg/ml), n=10**

RBx 10080758: 0.06

RBx 10115912: 0.5

Linezolid: 32

**Supplementary figure 1a**: *In vitro* activity of RBx 10080758 and RBx 10115912 against linezolid resistant (LZDres) *S. aureus.* RBx 10080758 and RBx 10115912 are active against LZDres *S. aureus.*

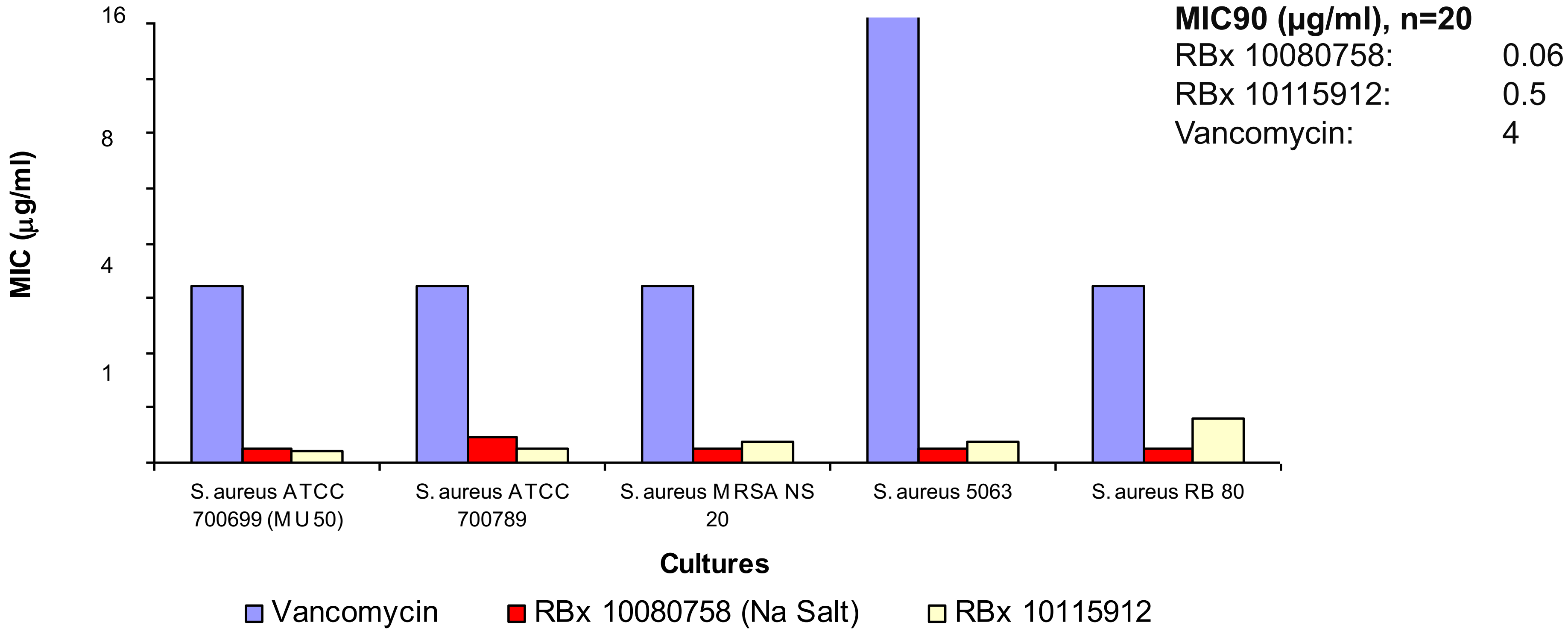


**Supplementary figure 1b**: *In vitro* activity of RBx 10080758 and RBx 10115912 against vancomycin resistant *S. aureus.* RBx 10080758 and RBx 10115912 are active against vancomycin intermediate *S. aureus* (VISA) and vancomycin resistant *S. aureus* (VRSA)

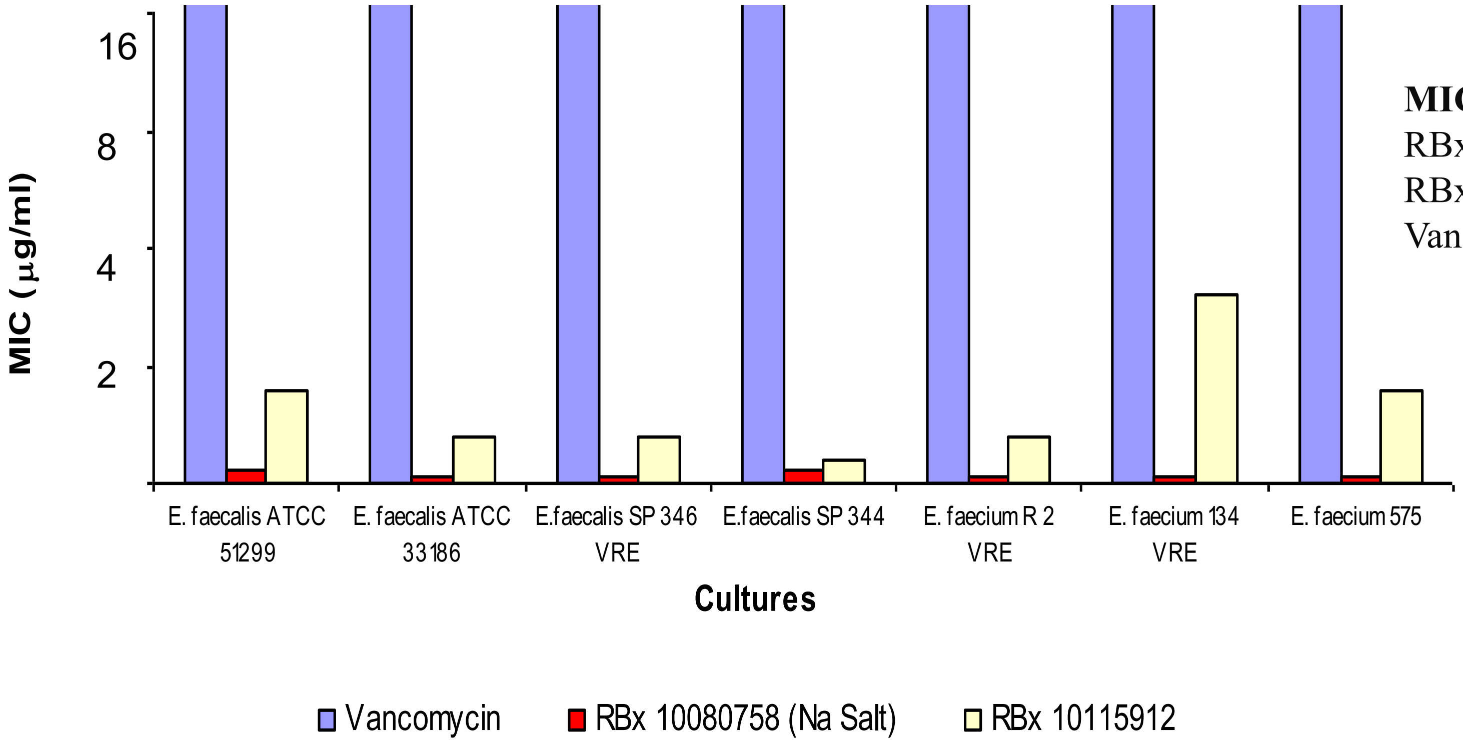


**MIC90 (µg/ml), n=20**

| | |
|---|---|
| RBx 10080758: | 0.06 |
| RBx 10115912: | 4 |
| Vancomycin: | 32 |

**Supplementary figure 1c:** *In vitro* activity of RBx 10080758 and RBx 10115912 against vancomycin resistant Enterococci (VRE). RBx 10080758 and RBx 10115912 are active against VRE

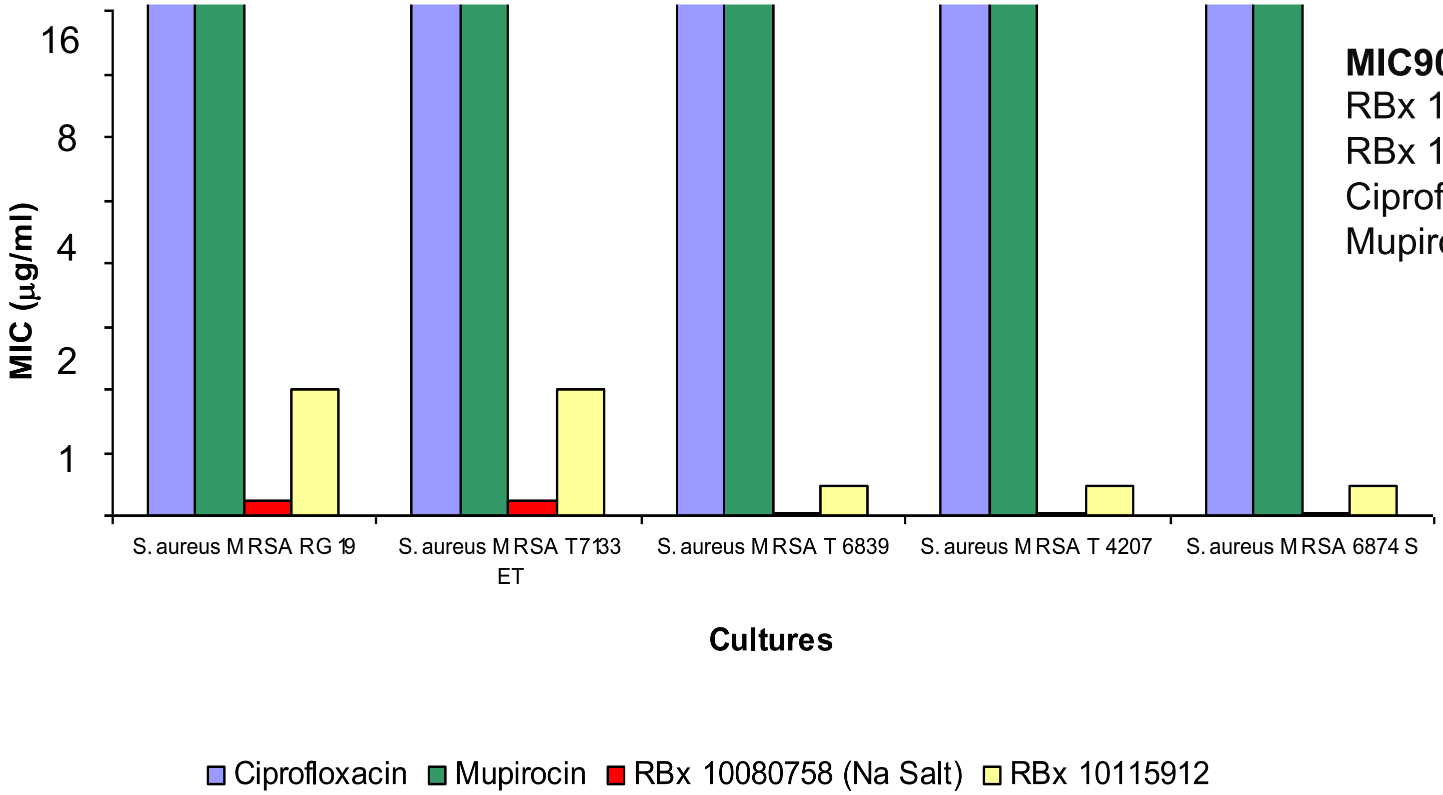


**MIC90 (µg/ml), n=20**

| | |
|---|---|
| RBx 10080758: | 0.06 |
| RBx 10115912: | 0.25 |
| Ciprofloxacin: | 16 |
| Mupirocin: | 512 |

**Supplementary figure 1d**: *In vitro* activity of RBx 10080758 and RBx 10115912 against multi-drug resistant *S. aureus.* RBx 10080758 and RBx 10115912 are active against MDR *S. aureus*

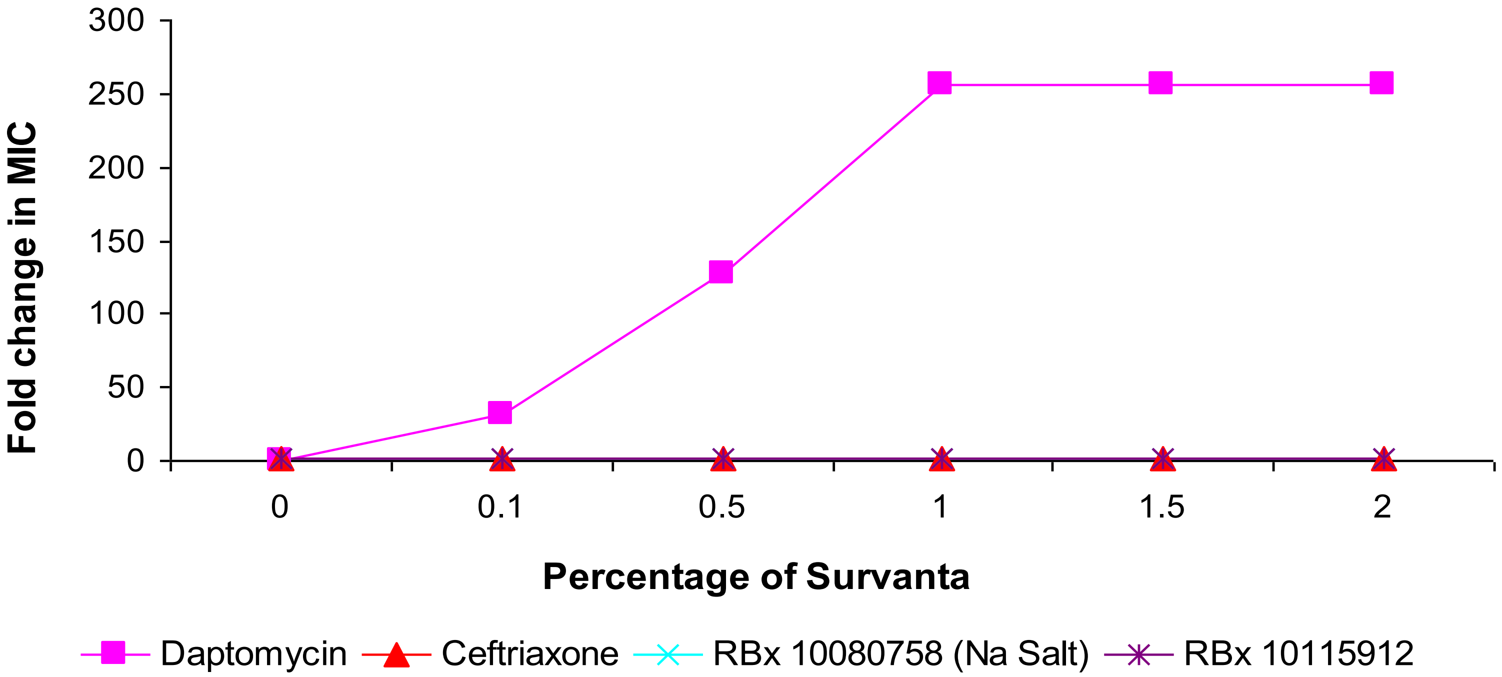


**Supplementary figure 2**: Effect of pulmonary surfactant on *in vitro* activity of RBx 10080758 and RBx 10115912 against *S. aureus* ATCC 29213. No effect on RBx 10080758 and RBx 10115912, however, daptomycin interact with pulmonary surfactant and inhibit antibacterial activity.
